# Isotopically Nonstationary ^13^C Metabolic Flux Analysis in Resting and Activated Human Platelets

**DOI:** 10.1101/2021.05.06.442995

**Authors:** Cara L. Sake, Alexander J. Metcalf, Jorge Di Paola, Keith B. Neeves, Nanette R. Boyle

**Affiliations:** Department of Chemical and Biological Engineering, Colorado School of Mines, Golden, CO, 80401 USA; Department of Pediatrics, Washington University School of Medicine, St. Louis, MO, 63110, USA; Department of Bioengineering, University of Colorado, Aurora, CO, 80045 USA; Hemophilia and Thrombosis Center, University of Colorado, Aurora, CO, 80045 USA; Department of Pediatrics, Section of Hematology, Oncology, and Bone Marrow Transplant University of Colorado, Aurora, CO, 80045 USA

**Keywords:** blood platelets, metabolic flux analysis, metabolomics, thrombin

## Abstract

Platelet metabolism is linked to platelet hyper- and hypoactivity in numerous human diseases. Developing a detailed understanding of the link between metabolic shifts and platelet activation state is integral to improving human health. Here, we show the first application of isotopically nonstationary ^13^C metabolic flux analysis to quantitatively measure carbon fluxes in both resting and thrombin activated platelets. Resting platelets primarily metabolize glucose to lactate via glycolysis, while acetate is oxidized to fuel the tricarboxylic acid cycle. Upon activation with thrombin, a potent platelet agonist, platelets increase their uptake of glucose 3-fold. This results in an absolute increase in flux throughout central metabolism, but when compared to resting platelets they redistribute carbon dramatically. Activated platelets decrease relative flux to the oxidative pentose phosphate pathway and TCA cycle from glucose and increase relative flux to lactate. These results provide the first report of reaction-level carbon fluxes in platelets and allow us to distinguish metabolic fluxes with much higher resolution than previous studies.

## INTRODUCTION

Platelets are anucleate cells that play a vital role in hemostasis, the physiological process of blood clot formation in response to vascular injury that occurs over the timescale of minutes. As such, much of what is known about platelet metabolism is derived from studies focused on signaling in response to acute exposure to agonists (Lepropre *et al*, 2018) and platelet interactions with other blood and vascular cells (Jaffe, 1987; Semple & Freedman, 2010). There have also been significant efforts in identifying the metabolic cause and effect of platelet storage lesion in transfusion medicine (Scott, 1995; Scott *et al*, 1992). Beyond hemostasis, platelets are increasingly recognized for their various roles in innate immunity, diabetes, hyperglycemia, atherosclerosis, kidney disease, and coronary artery disease (Burkhart *et al*, 2014; Kramer, 2014; van der Meijden & Heemskerk, 2019). Recent studies in these chronic disorders suggest that changes in platelet metabolism play a role in their pathophysiology and can potentially be used as a biomarker (Davizon-Castillo *et al*, 2019; McDowell *et al*, 2020).

Previous studies on platelet metabolism have deduced metabolic pathway trends as a result of storage (Paglia *et al*, 2014), old age (Davizon-Castillo *et al.*, 2019), activation (Ravi *et al*, 2015), and in response to specific reaction or pathway inhibitors (Ju *et al*, 2012). Primarily, these studies have monitored metabolic markers that indicate changes in whole metabolic pathways. Such techniques include radiometric labels (Cohen & Wittels, 1970), respirometry (Sowton *et al*, 2018), and extracellular flux analysis (Fidler *et al*, 2017). For example, Seahorse extracellular flux (XF) analyzer measures oxygen consumption rate (OCR) and extracellular acidification rate (ECAR), which are interpreted as indirect measurements of mitochondrial oxidative phosphorylation and glycolysis, respectively (Nayak *et al*, 2018). While informative, these measurements can be challenging to interpret, in part due to the existence of multiple acidification mechanisms and the technique’s reliance on sequentially administered inhibitors to probe the cellular utilization of glucose (Kholmukhamedov & Jobe, 2019). Metabolic pathways are complex, and attenuation of an individual reaction within a pathway does not necessarily result in attenuation of the whole pathway, further complicating the pursuit of interpreting extracellular flux measurements.

Metabolomics has been performed extensively on platelets during storage (Jóhannsson *et al*, 2018; Paglia *et al*, 2015; Paglia *et al.*, 2014) and after chemical treatment (Abonnenc *et al*, 2016; Ju *et al.*, 2012) to identify changes in metabolite pool sizes. While this technique can provide a global view of changes in metabolism, *how* metabolite pool sizes change remains unknown. Quantitative measurement of reaction fluxes is needed to identify how metabolism was rewired to lead to the changes in metabolite concentration. Isotope assisted metabolic flux analysis (^13^C-MFA) is the current gold standard to quantitatively measure carbon fluxes in a cell by tracking a stable isotopic tracer, in this case ^13^C, through metabolism (Wiechert, 2001). Experimental data is combined with computational metabolic models to deconvolute the data and determine intracellular fluxes. Enzymes rearrange the atoms in metabolites, and ^13^C-MFA uses isotope labels to measure the rearrangements of carbon atoms; these labeling patterns, or isotopomers, can then be mathematically fit with established stoichiometric metabolic models and known atom transitions to obtain intracellular fluxes (Cheah & Young, 2018). This technique provides experimentally derived fluxes, as well as the ability to validate metabolic network models that include compartmentalization, parallel pathways, and cyclic reaction sets (Antoniewicz, 2015) and has been successfully implemented to measure fluxes in a variety of organisms including bacteria, plants and mammalian cells.

Platelets present unique challenges for developing metabolic models and obtaining experimental ^13^C-MFA measurements. First, unlike other cells that ^13^C-MFA has been applied to, platelets are anuclear. This means they do not replicate and are unable to respond to environmental cues by altering mRNA abundance. Moreover, their lifetime is limited to 7-10 days both inside and outside of the body (Quach *et al*, 2018). Additionally, platelets are known to display unique phenotypes across storage lengths (Paglia *et al.*, 2014), which potentially limit the timescale for experimental measurements, especially during steady state labeling experiments, which often require incubation with isotope tracers for several days (Dai & Locasale, 2016).

In this work, we describe the *first* application of isotopically nonstationary metabolic flux analysis (INST-MFA) to measure carbon fluxes in resting and activated platelets. To our knowledge, this is the first report using ^13^C-MFA to measure metabolic flux in an anuclear cell. We establish metabolic steady state in platelets, define a metabolic model to track atom transitions throughout platelet central metabolism, and *quantitatively measure* metabolic fluxes of resting and thrombin activated platelets.

## RESULTS

### Metabolic modeling suggests glucose and acetate tracers

The workflow for implementing INST-MFA in platelets requires experimental, analytical, and computational approaches (Figure 1). It is common to start on the computational model, which can aid in experimental design. The metabolic network of platelet central metabolism used is shown in Figure 2 and was constructed in the isotopomer network compartmental analysis (INCA) software, a MATLAB-based software package for INST-MFA (Young, 2014). The experimental tracers used for ^13^C-MFA determines the confidence of calculated flux results, therefore care was taken during this step to select a mixture that produced unique transient labeling behavior and several isotopomers per metabolite that remained enriched over the course of the experiment. Constraints derived from the iAT-PLT-636 model (Thomas *et al*, 2015) were used with the INCA *tracer simulation* function to identify glucose and acetate as viable carbon sources for probing the glycolysis/pentose phosphate pathway and the tricarboxylic acid (TCA) cycle, respectively. Acetate is an oxidative fuel that enters the TCA cycle via acetyl-CoA (Gulliksson, 1993; Guppy *et al*, 1990). Simulations with a variety of readily available isotopic tracers revealed that mixtures of [1,2-^13^C_2_]glucose, [U-^13^C_6_]glucose, [1-^13^C]acetate, [2-^13^C]acetate, and their unlabeled counterparts provided favorable steady state labeling distributions (Supplemental Fig. 1).

**Figure 1.**
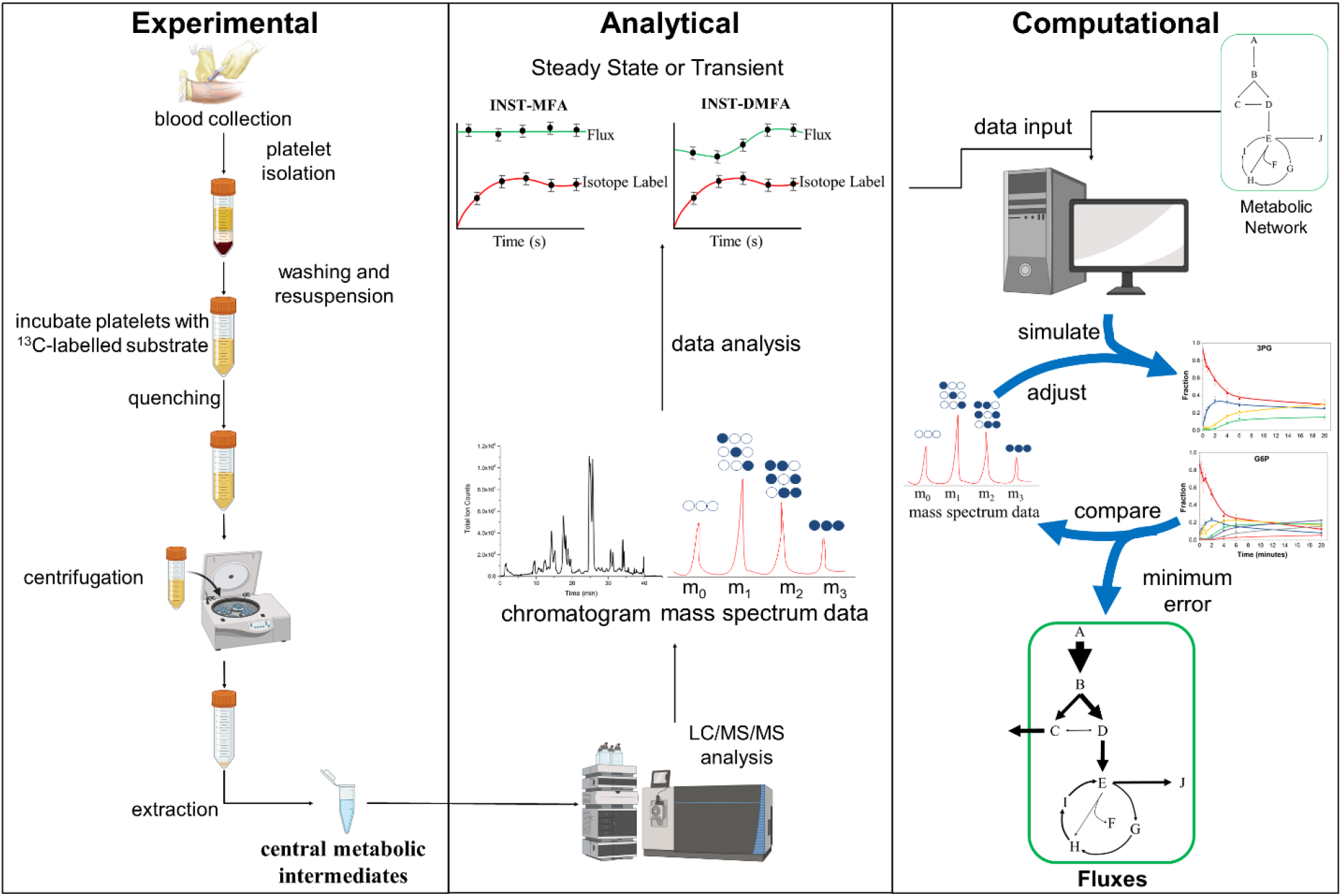
Summary of the workflow for performing ^13^C-MFA in platelets. Washed platelets isolated from whole blood are incubated with ^13^C-glucose or ^13^C-acetate and then rapidly sampled, quenched, and intracellular metabolites extracted. The metabolite profile is analyzed using LC-MS/MS to obtain mass isotopomer distributions (MIDs). The analytical data is then paired with the modeled reaction network and respective atom transitions; fluxes are determined through iterative parameter adjustment to minimize the error between the simulated and measured isotopomer profiles. Figure adapted from (Sake *et al.*, 2019).

**Figure 2.**
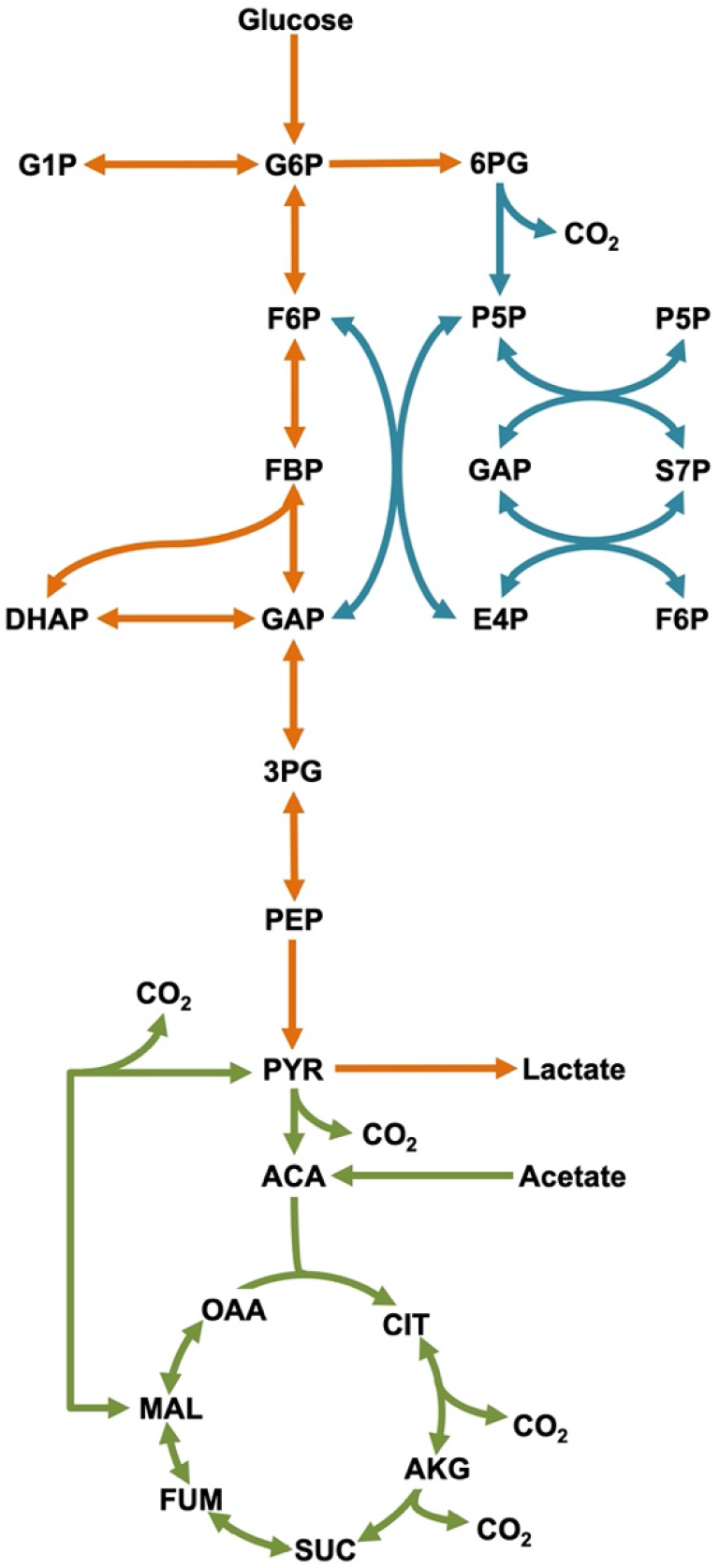
Platelet central metabolic network. Reactions that are reversible are shown with two-headed arrows. Metabolic pathways are color coded to represent glycolysis (orange), pentose phosphate pathway (blue), and the tricarboxylic acid (TCA) cycle (green).

### Washed platelets maintain short-term metabolic steady state

A key assumption of ^13^C-MFA is that cellular metabolism remains approximately at steady state throughout the course of the labeling experiment (Noh *et al*, 2006), which occurs when the pool size of metabolites remains unchanging with time. For conventional metabolomic studies in bacteria, metabolic steady state is assumed while the culture is in the exponential phase of growth (Noh *et al.*, 2006). Similarly for mammalian cell studies, steady state can typically be assumed for continuous cultures (Wiechert & Noh, 2005). To confirm metabolic steady state is a reasonable assumption over the timescale of our experiments, we tracked the total pool size of each targeted metabolite for one hour following washing and a 90-minute resting period. Representative metabolites from resting platelets with a vehicle control and platelets activated with 1 U/mL thrombin are shown in Figure 3. Separate studies of metabolite pool size in washed platelets across multiple donors showed similar results (Supplemental Fig. 2).

**Figure 3.**
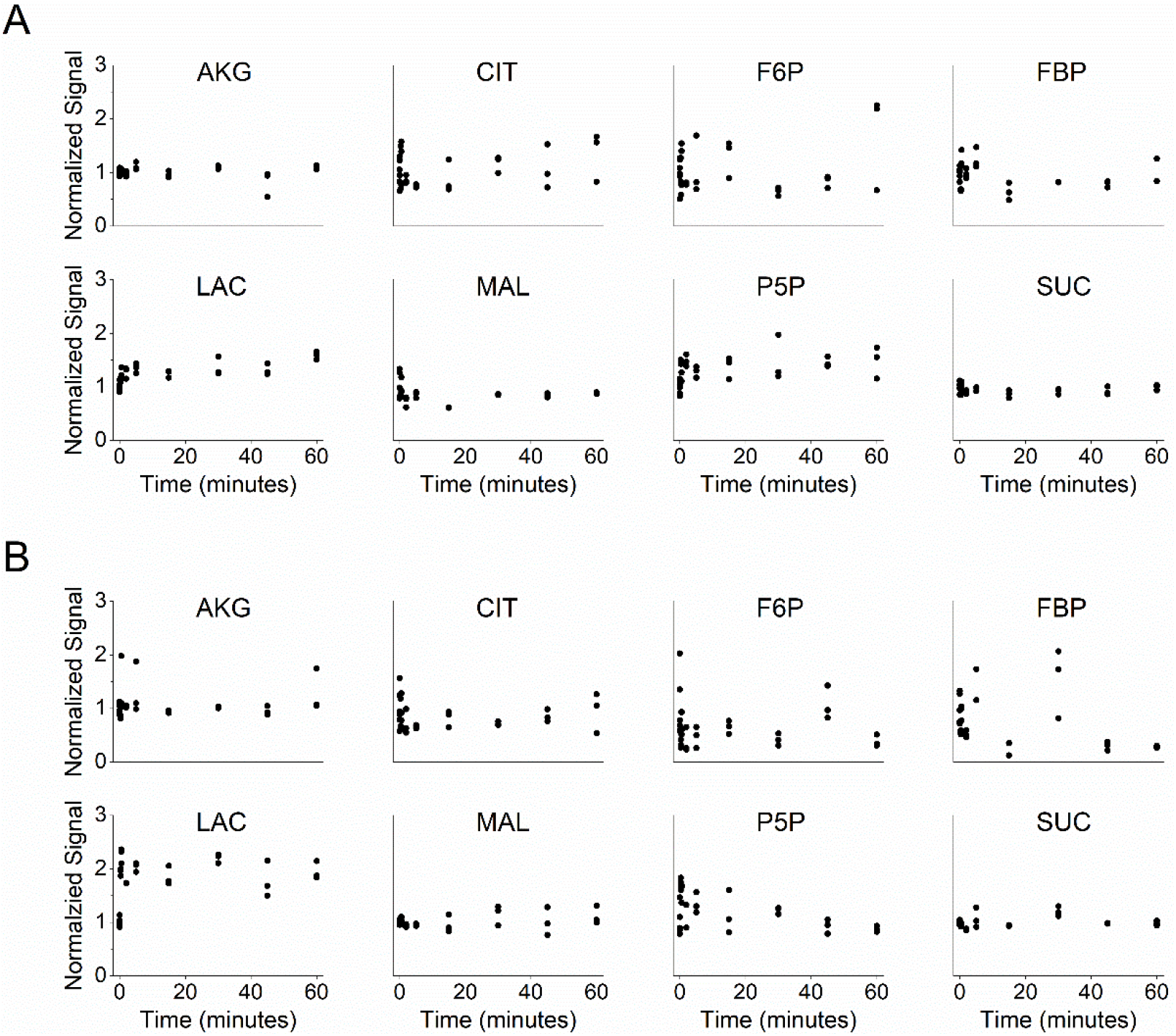
Metabolite pool size of representative metabolites over 60 minutes. Pool size for (A) resting (vehicle = ethanol) and (B) thrombin activated (1 U/mL) platelets is represented by the sum of the MS signal for each metabolite’s isotopomer, displayed normalized to the mean at time = 0 min. See also Supplemental Fig. 2.

### Isotope labeling dynamics and extracellular flux measurements

Uptake and excretion rates are parameters that constrain ^13^C-MFA models. We measured the uptake of both carbon sources, glucose and acetate, as well as the production of lactate over the course of our experiments (Figure 4). When treated with 1 U/mL thrombin, washed platelets increased overall glucose consumption (exogenous glucose and stored glycogen) by 4.2-fold and lactate production by 3.2-fold, suggesting that the increase in glucose consumption fuels additional pathways outside glycolysis and lactate production. Acetate uptake remained stable with and without treatment with thrombin, and on a carbon atom basis, was 84% of the glucose uptake of resting platelets. These uptake and excretion fluxes were used to constrain the INST-MFA model alongside isotopic labeling measurements.

**Figure 4.**
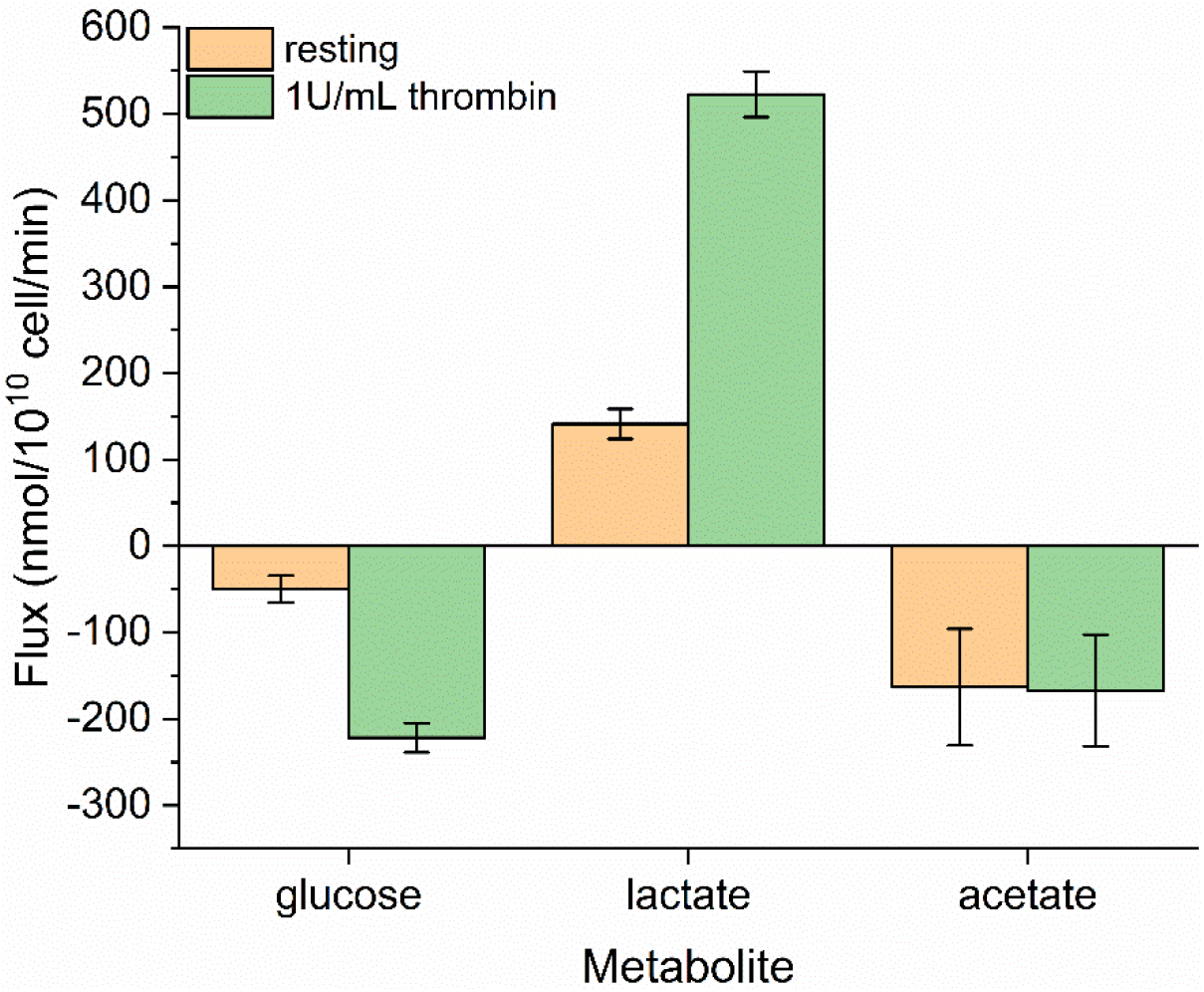
Measured uptake and excretion fluxes for resting (orange) and thrombin activated (green) platelets. Glucose, lactate, and acetate fluxes are shown as negative for uptake/carbon source and positive for excretion/carbon sink. Extracellular metabolite concentration was measured from the reserved carbon labeling experiment supernatants. The reported flux represents the slope of a linear regression to the measured extracellular concentrations using a least squares method. Error bars represent standard error.

To further probe platelet metabolism, parallel labeling experiments were performed with glucose as the primary tracer for labeling glycolysis and the pentose phosphate pathway. Glutamine is suggested as an optimal tracer for probing the TCA cycle of mammalian cells (Metallo *et al*, 2009) and has been used successfully in parallel labeling experiments (Ahn & Antoniewicz, 2013); however there is uncertainty regarding whether glutamine metabolism in platelets is complete (Scott *et al.*, 1992). We ruled out glutamine as a tracer after experimental trials with [U-^13^C_5_]glutamine showed poor enrichment and slow labeling dynamics in TCA metabolites (Supplemental Fig. 3), and instead we opted for acetate as the secondary tracer. The dynamic glucose and acetate labeling measurements and INST-MFA model fits of select metabolites representing all three major central metabolism pathways are shown in Figure 5. Fits for remaining metabolites are shown in Supplemental Figs. 4-6. The data do not indicate significant differences in labeling speed during thrombin activation but do show a higher overall ^13^C-enrichment (total fraction of carbon enriched with ^13^C) for glycolysis metabolites.

**Figure 5.**
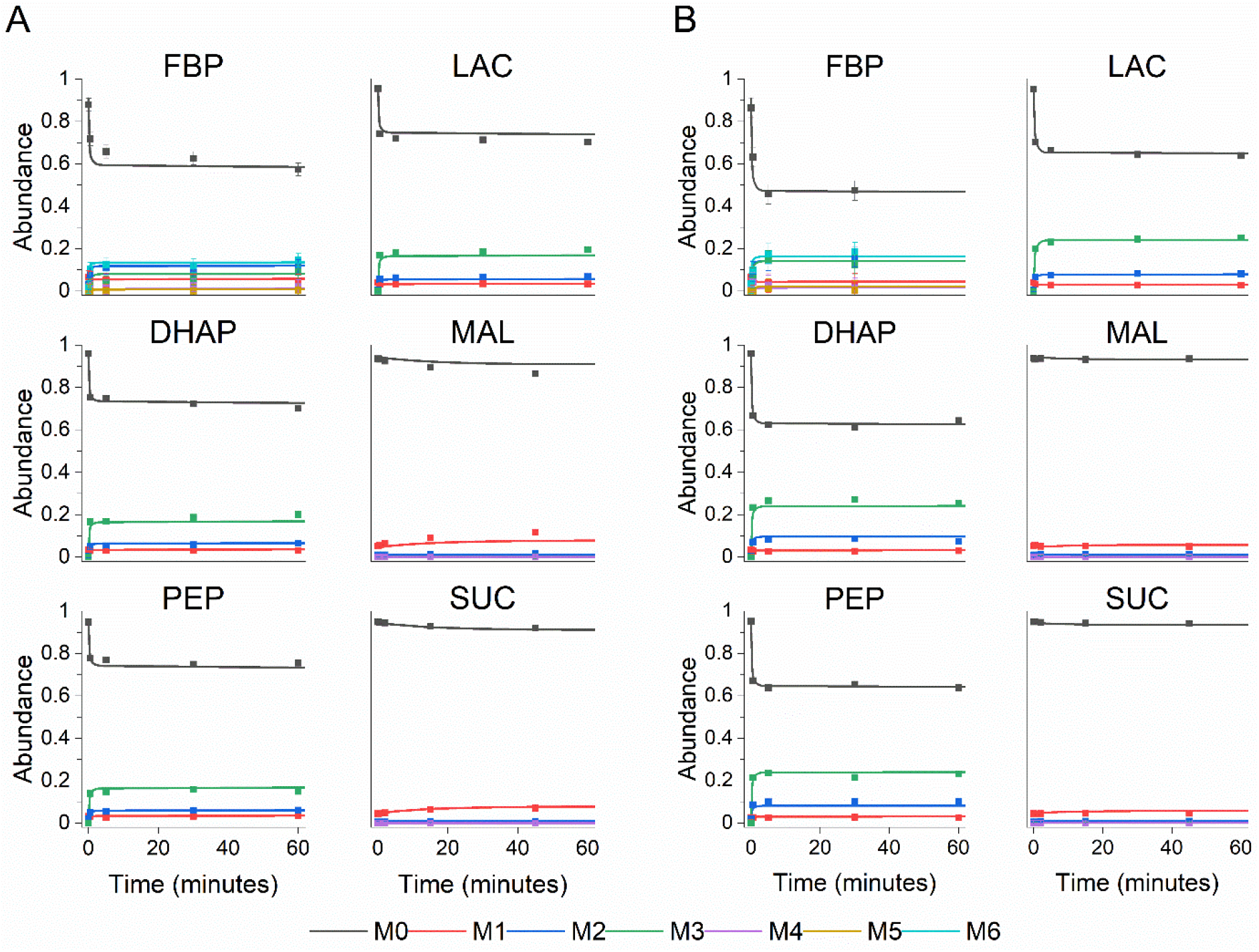
Experimentally measured mass isotopomer abundances (symbols) and INST-MFA model fits (lines). Each panel shows the ^13^C labeling trajectories as fractional abundance of fructose bisphosphate (FBP), dihydroxyacetone phosphate (DHAP), phosphoenolpyruvate (PEP), lactate (LAC), malate (MAL), and succinate (SUC) for (A) resting and (B) thrombin activated platelets. Labeling for FBP, DHAP, PEP, and LAC originate from the glucose tracer and labeling for MAL and SUC come from the acetate tracer. Raw mass isotopomer abundances are shown without correction for natural abundance. Error bars represent standard measurement error.

### Metabolic flux analysis does not indicate significant amino acid metabolism

In addition to central carbon metabolites, labeling profiles were measured for four amino acids: alanine, aspartate, glutamine, and glutamate (Supplemental Figs. 4-6). Alanine and glutamine showed enrichment close to zero in all experiments. The labeling profile of glutamate was similar to that of malate. Aspartate was enriched from the acetate tracer during the resting experiment but was enriched from both glucose and acetate during the thrombin experiment (Supplemental Fig. 6). Aspartate labeling proved particularly useful for verifying the fate of glucose, as ^13^C-enrichment in aspartate rose to 5% with added thrombin when glucose was the tracer (Supplemental Fig. 6).

**Figure 6.**
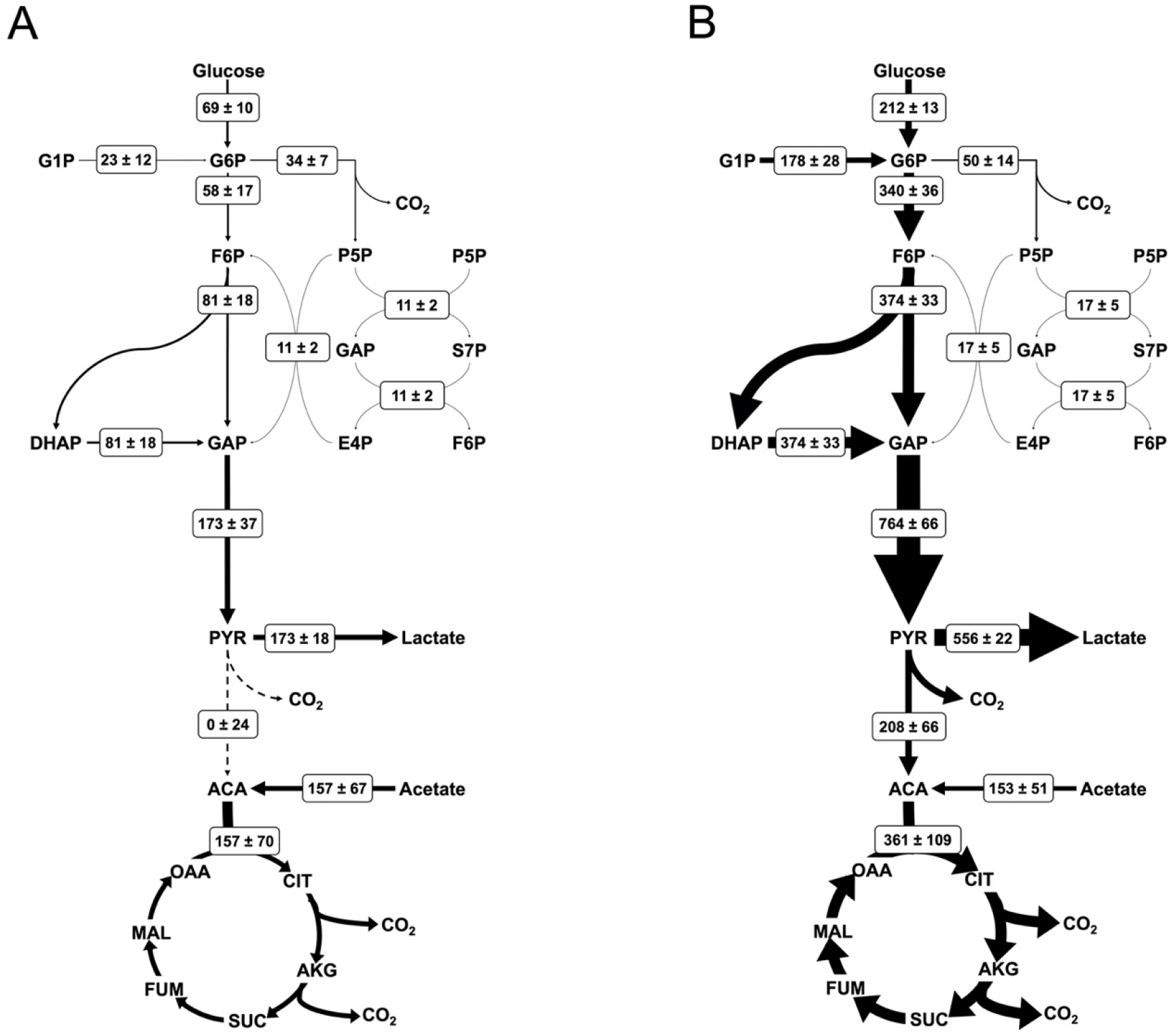
Flux maps of platelet metabolism. Net fluxes are shown for (A) resting and (B) thrombin activated conditions in the form M±SE where M is the calculated flux and SE is the standard error of the 95% confidence interval between upper and lower bounds with units of nmol/10^10^ platelets/min. Arrow thickness is linearly proportional to the net flux, with dashed lines representing zero net flux. All calculated net fluxes and their 95% confidence interval are listed in Supplemental Table 1.

### Metabolic flux analysis reveals glycolytic rerouting upon thrombin stimulation

Mass isotopomer abundances for 25 metabolites and six extracellular flux measurements (three for each parallel labeling experiment) were used to fit the INST-MFA model for each condition. Both resting and thrombin conditions were fitted with the sum-of-squared residuals between computational and experimental measurements falling within the expected ranges for a 95% confidence interval; SSR = 610.7 with range [518.0 651.8] for vehicle control and SSR = 525.5 with range [448.4 573.4] for thrombin. Measured metabolic fluxes for resting and thrombin activated platelets are presented in Figure 6. Washed, resting platelets in the presence of glucose and acetate display balanced, yet fragmented metabolism between glycolysis and the TCA cycle. Glucose is stoichiometrically converted to lactate via glycolysis. A significant portion of glucose metabolism (37% of the flux through G6P) is directed through the oxidative pentose phosphate pathway. Our results show that acetate is the preferred substrate for oxidative metabolism in resting platelets, not glucose.

Upon activation with thrombin, platelet metabolism undergoes global increases. Flux through the oxidative pentose phosphate pathway increases by 1.4-fold and TCA cycle flux increases by 2.3-fold. The greatest augmentation occurs through glycolysis, where the flux increases 4.4-fold, resulting in significant increases in lactate production. This follows the trends observed with increased glucose uptake and lactate excretion in thrombin activated platelets (Figure 4). Importantly, thrombin activation causes an absolute increase in glucose oxidation via the TCA cycle (Figure 6). Interestingly, when fluxes are normalized to resting platelets, the relative flux through the TCA cycle actually decreases by approximately 25% (Figure 7). All major CO_2_ producing pathways show an increase in flux with thrombin stimulation leading to an overall 2.8-fold increase in CO_2_ production.

**Figure 7.**
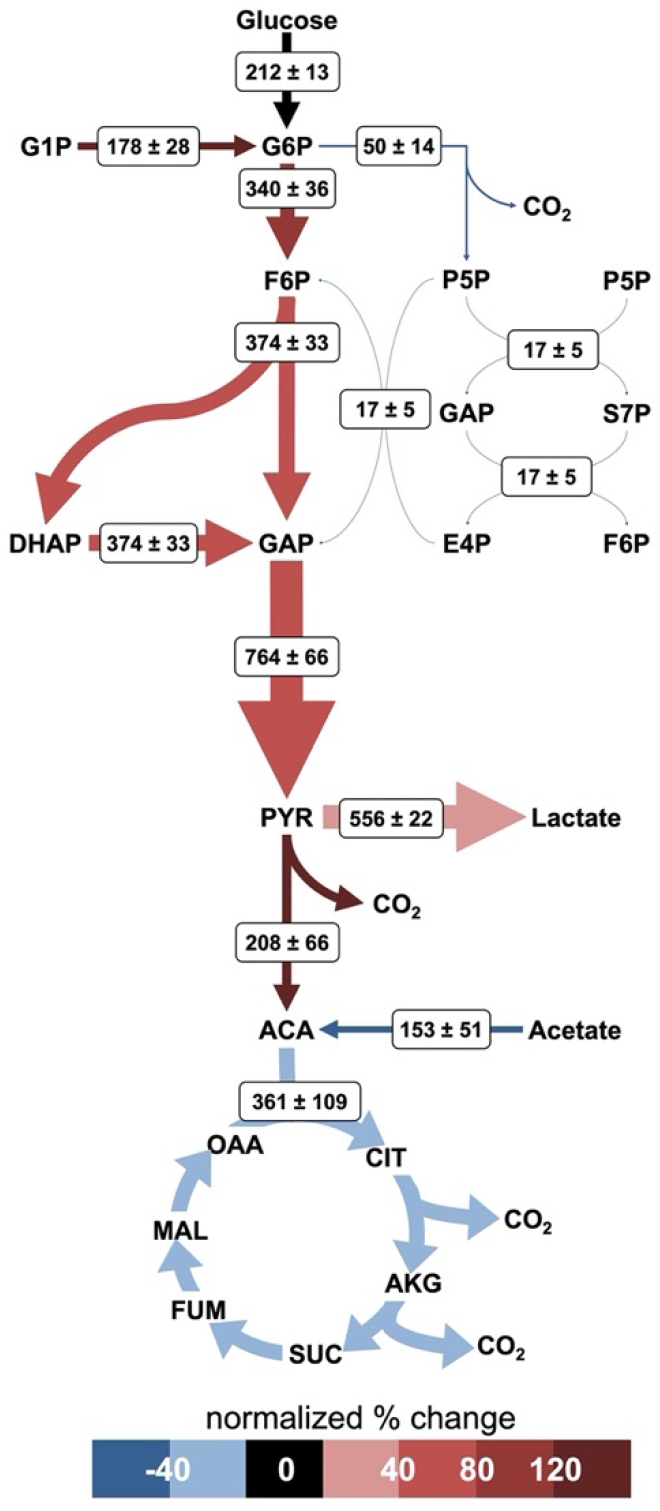
Normalized fluxes of activated platelet metabolism. Net fluxes and arrow thicknesses are shown as previously described for Fig 6. Arrow color represents the percent change of the thrombin condition normalized flux from the resting condition normalized flux. Fluxes are normalized to the total glucose uptake rate.

## DISCUSSION

The goal of this study was to quantitatively measure central metabolic fluxes in resting and thrombin activated human platelets using ^13^C-MFA. In order to use ^13^C-MFA for the first time on human platelets, we had to address the unique challenges posed by platelet physiology. Although previous studies have measured platelet metabolism through global and indirect methods, and even proposed predicted platelet flux maps (Jóhannsson *et al.*, 2018; Paglia *et al.*, 2014; Thomas *et al.*, 2015), these techniques lack the reaction-level resolution and direct quantitative solutions provided by MFA. The approach taken here offers the primary advantage of removing inference in interpreting results: carbon flux measurements are quantitative, include upper and lower bounds, and measurements are not subject to confounding factors that are present in more global measurement types.

Resting washed human platelets experience balanced yet divided metabolism in which glucose consumption fully fuels lactate production from glycolysis and acetate uptake solely fuels the TCA cycle. Glycolysis and TCA flux are approximately equal, with TCA flux being >90% of glycolytic flux. A significant portion of G6P carbon (37%) enters the pentose phosphate pathway. Thrombin activated platelets experience global increases in intracellular pathway fluxes, fueled by a 3-fold increase in glucose uptake, without any changes in acetate uptake. As a consequence, increases to oxidative mitochondrial metabolism require a metabolic shift, in which the previously unused pyruvate oxidation pathway is activated to redirect 27% of the glycolytic flux to the TCA cycle. Thrombin activated platelets also display increases in pentose phosphate pathway behavior, for a 1.4-fold change from the resting case; yet proportionally, thrombin treatment lowers pentose phosphate flux to 13% of the total glycolytic flux.

INST-MFA calculated fluxes for resting platelets describe a central metabolism in which glucose is fully metabolized to lactate through glycolysis with approximately equal flux activity in the TCA cycle. In line with our results, many previous studies have observed stoichiometric conversion of glucose to lactate in resting platelets (Guppy *et al.*, 1990; Scott *et al.*, 1992; Vasta *et al*, 1993). While originally thought to rely solely on oxidative phosphorylation, resting platelets are known to utilize both glycolysis and oxidative phosphorylation (Chacko *et al*, 2013; Nayak *et al.*, 2018). Untargeted metabolomics of resting platelets suggest that platelet metabolism follows unique phenotypes related to storage length (Paglia *et al.*, 2014); because our study was performed within hours post-collection from whole blood, the platelets measured here are expected to be comparable with the short-term storage phenotype associated with active glycolysis, active pentose phosphate pathway, and down-regulated TCA cycle (Paglia *et al.*, 2014). Our results do support this summary, showing the greatest flux through glycolysis at 173 ± 37 nmol/10^10^ platelets/min, with 37% of G6P flux entering the pentose phosphate pathway, and the TCA flux equaling 90% that of glycolysis at 157 ± 70 nmol/10^10^ platelets/min.

Platelet activation with thrombin has been reported to cause significant changes to platelet metabolism, particularly in regard to glucose uptake, extracellular acidification, and oxygen consumption (Aibibula *et al*, 2018; Nayak *et al.*, 2018; Ravi *et al.*, 2015). Agonist induced platelet activation initiates a step change in energy requirements, and as a result, activated platelets exhibit increased oxidative phosphorylation and glycolysis to accommodate increased ATP demands (Kholmukhamedov & Jobe, 2019). Previous studies have observed that activated platelets show increased glycolysis relative to mitochondrial phosphorylation (Kramer, 2014). Often this behavior of thrombin activated platelets is described as a “glycolytic phenotype” (Aibibula *et al.*, 2018; Kholmukhamedov & Jobe, 2019). Our results support these prior studies, as we see the largest fold-changes from the resting to activated state occur in glycolysis.

Using INST-MFA we now have a finer grain understanding of the platelet glycolytic phenotype, wherein a dual increase in glucose oxidation and fermentative glycolysis (sometimes referred to as *aerobic glycolysis* in other works to describe lactate production in the presence of oxygen) is observed following thrombin stimulation. The glycolytic flux increases 4.4-fold compared to the resting platelet condition, 73% of which contributes to a 3.2-fold increase in lactate production with the balance fueling the previously unused pathway for pyruvate oxidation. We measured universal increases in absolute flux throughout all major pathways: glycolysis, pentose phosphate pathway, and the TCA cycle. However, we also observed a metabolic switch in which resting platelets perform no glucose oxidation, instead relying on acetate to feed TCA metabolism, and thrombin activated platelets incur large increases in glucose uptake in order to support increased fermentative glycolysis *and* glucose oxidation via the TCA cycle. For this reason, it is more appropriate to also describe resting platelets as displaying a glycolytic phenotype, because their utilization of glucose is 100% glycolytic and 0% oxidative, a state in which glucose is stoichiometrically converted to lactate.

Thrombin activated platelets do display a glycolysis dominant phenotype in terms of absolute flux. However, from the perspective of glucose utilization, 27% of the glycolytic flux in thrombin activated platelets is oxidized in the TCA cycle, meaning that thrombin activated platelets trend toward a more oxidative phenotype than resting platelets. Acetate uptake remains unchanged between the two cases, allowing overall increases in TCA cycle flux as carbon from glucose and acetate uptakes are combined at the first step of the TCA cycle with acetyl-CoA.

In this study, we were also able to further define mitochondrial activity in terms of TCA cycle flux and quantify the magnitude between the two pathways. Resting platelets rely on glycolysis and mitochondrial TCA cycle approximately equally, while platelets activated with thrombin preferentially metabolize glucose for 4.4-fold increases in glycolysis, relative to 2.3-fold increases throughout the TCA cycle.

Extracellular fluxes were measured from major carbon sources and sinks: glucose, lactate, and acetate. CO_2_ is also a carbon sink, primarily from the TCA cycle, but there is also a non-trivial contribution to CO_2_ production from the oxidative pentose phosphate pathway. A unified biomass equation was purposefully neglected on the basis that *in vivo*, platelets arise via differentiation from megakaryocytes and therefore do not divide outside the body. Precedent shows that neglecting the biomass equation is suitable during slow cell growth (Egnatchik *et al*, 2014): a lack of platelet division indicates that platelet growth rate is negligible. As a consequence of their origin, platelets lack a nucleus; however, they do contain mRNA and therefore can synthesize protein (Weyrich *et al*, 2009). The possibility of net flux leaving central metabolism to support protein synthesis was considered but was ultimately rejected. Again, there is precedent for neglecting the contribution of protein turnover, especially during INST-MFA when the labeling time of metabolic intermediates is expected to be significantly shorter than that of proteins (Schaub *et al*, 2008). However, to account for the possibility, alanine, aspartate, glutamate, and glutamine were included in the metabolic network. Flux results showed no net flux entering the amino acid pools, but their labeling profiles remained useful for flux calculation via exchange flux (see Supplemental Table 2).

A limitation of this study is the use of washed platelets. Cellular behavior is dependent on extracellular conditions, especially the availability of various carbon sources. For the purpose of isotope labeling, we choose to use washed platelets to avoid glucose carryover, which would require quantification of the dilution of the isotope tracer. However, we recognize that this deviates from the in vivo environment of circulating platelets. Further studies are needed to examine how platelet metabolism varies in more physiologic media like plasma or whole blood.

In summary, we have applied ^13^C-MFA for the first time in platelet cells to measure the central metabolism fluxes in resting and thrombin activated platelets. Increased understanding of platelet metabolism has the potential to guide platelet storage, serve as a biomarker for platelet phenotype, and provide a new tool of mechanistic studies of platelet physiology. Our model has the potential to be expanded to include secondary pathways linked to platelet activation including fatty acid (Ravi *et al.*, 2015; Slatter *et al*, 2016) and eicosanoid (Chen & Silverstein, 2018; Ju *et al.*, 2012) metabolism. Such pursuits may require alternative carbon labeling sources such as palmitate or arachidonic acid to successfully probe downstream pathways. Certain assumptions, such as metabolic steady state, may need to be reassessed before adapting to other conditions; if the metabolic steady state condition cannot be upheld, other specialized ^13^C-MFA techniques such as dynamic metabolic flux analysis (Antoniewicz, 2013) may be necessary.

## Materials and Methods

A comprehensive list of abbreviations is available in Supplemental Table 3.

### Ethics approval and consent to participate

The study received Institutional Review Board approval from the University of Colorado Anschutz Medical Campus.

### Platelet collection and isolation

Blood samples were collected by venipuncture using a 21-gauge needle and collected into a syringe containing sodium citrate (3.8%). Anticoagulant citrate dextrose (ACD) solution was added to the whole blood in a 1:10 ratio, and the mixture was centrifuged at 200*g* to fractionate. Platelets were isolated and washed from the platelet rich plasma (PRP). In short, platelets were washed in modified Tyrode’s buffer with prostacyclin (PGI2) as an anticoagulant. Platelets were pelleted at 1000*g* for 10 minutes. After washing, isolated platelets were resuspended in Tyrode’s buffer, counted, and adjusted to a concentration of 8×10^8^ cell/mL. Platelets were incubated at 37°C for 90 minutes prior to performing experiments.

### Carbon labeling experiment

Prior to labeling, platelet suspensions were incubated with vehicle (ethanol) or thrombin (1 U/mL) for 15 minutes. Glucose and acetate tracers were administered in parallel labeling experiments and samples were taken at predetermined intervals over 60 minutes. Added glucose was composed of 30% [1,2-^13^C_2_]glucose, 50% [U-^13^C_6_]glucose, and 20% unlabeled glucose. Added acetate was composed of 25% [1-^13^C]acetate, 25% [2-^13^C]acetate, and 50% unlabeled acetate. Samples were collected in triplicate at 0, 30 s, 5 min, 30 min, 60 min, and 180 min for glucose labeling and 0, 20 s, 2 min, 15 min, 45 min, and 180 min for acetate labeling. The 0 time-point samples were taken immediately prior to the addition of the label. Samples were quenched by rapidly plunging 0.5 mL portions into 1 mL of pre-chilled partially frozen normal saline and centrifuged to pellet at 1600*g* and 0°C. The supernatant was collected for extracellular metabolite analysis and the pellet was stored at −20°C prior to extraction.

### Extracellular flux measurements

Glucose uptake, lactate excretion, and acetate uptake rates were determined by analyzing the change in the metabolites’ concentrations from the saved supernatants of the labeling experiment. The retained supernatant was thawed, mixed well, and a portion was removed for each assay. For glucose, the supernatant was analyzed using the Sigma starch assay kit (SA20). The lactate concentration was determined using the Megazyme L-lactic assay kit (K-LATE). Acetate was analyzed using the BioAssay EnzyChrom acetate assay kit. Concentration data over time was linearly fit using a least squares method, where the slope and its standard error were taken as extracellular flux.

### Metabolite extraction

Intracellular metabolites were extracted using a method adapted from (Paglia *et al*, 2012). Platelet pellets were resuspended in 500 μL of 70% methanol and spiked with ribitol and PIPES as internal standards. Samples were frozen in liquid nitrogen, thawed on ice, and vortexed at 0°C for 5 minutes and 1000 rpm. This freeze/thaw/vortex cycle was repeated twice more, and samples were centrifuged at 16,000g for 5 minutes at −4°C in a Sorvall Legend Micro 17R microcentrifuge (Thermo Scientific). The supernatant (extract) was collected in a new tube and stored at −20°C. The extraction process was then repeated with 500 μL of 50% methanol and the second collected supernatant was added to the first. The samples were then dried overnight under vacuum in a Savant SPD131DDA SpeedVac (Thermo Scientific). Dried extracts were resuspended in 150 μL Optima water and filtered with nylon filter tubes (Spin-X, Costar). Filters were rinsed with an additional 50 μL water for a total final extract volume of 200 μL for LC-MS/MS analysis.

### Analytical methods

Metabolite extracts were analyzed for targeted metabolites listed in Supplemental Table 4 using an LC-MS/MS method adapted from (Young *et al*, 2011). A Phenomenex 150 mm × 2 mm Synergi Hydro-RP column was used on an Agilent 1200 Series HPLC system coupled to an AB Sciex 5500 QTRAP system. LC was performed with an injection volume of 20 μL, using gradient elution of 10 mM tributylamine and 15 mM acetic acid (aqueous phase) with acetonitrile (organic phase) at a constant flow of 0.3 mL/min. The gradient profile of the organic phase is as follows: 0% B (0 min), 8% B (10 min), 16% B (15 min), 30% B (16.5 min), 30% B (19 min), 90% B (21.5 min), 90% B (26.5 min), 0% B (26.6 min), and 0% B (30.5 min). MS analysis was performed in negative mode using a multiple reaction monitoring (MRM) acquisition method. Data acquisition was performed on the ABSciex Analyst 1.7 software. Absolute quantification of intracellular metabolites was performed using the Analyst MultiQuant 3.0.3 Software. For isotope labeling profiles, MSConvert was used to convert LC-MS/MS data files (ABSciex *.wiff* and *.wiff.scan*) to an open-source format (Holman *et al*, 2014). A combination of the pyOpenMS (Rost *et al*, 2014) and SciPy (Virtanen *et al*, 2020) packages in Python, was used for the process of peak finding, window sizing, and noise assessment across multiple isotopes.

### Isotopomer network and flux calculations

An isotopomer network model describing human platelet central metabolism was constructed primarily from the published, manually curated reaction network iAT-PLT-636 (Thomas *et al.*, 2015). Carbon atom transitions were assigned with the assistance of the carbon fate maps from (Mu *et al*, 2007). Isotopic nonstationary metabolic flux analysis (INST-MFA) was performed in the Isotopomer Network Compartmental Analysis (INCA) software, along with the corresponding least-squares parameter regression and sensitivity analysis (Young, 2014). A list of the reactions and atom transitions for the model is listed in Supplemental Table 5.

## Acknowledgements

Thank you to Allaura Cox from the Di Paola group for kindly sharing her platelet washing protocol.

## Funding

This project is supported by the National Heart, Lung, and Blood Institute of the National Institutes of Health under award number R61HL141794.

## Author contributions

NB, KN, JDP, and CS conceived the project. CS performed statistical and experimental analysis for the steady state pool size study; data collection, analysis, and modeling for ^13^C-MFA; and was the primary contributor in writing the manuscript. AM supported construction of the reaction network, ran labeling simulations, built the code utilized for LC-MS/MS data analysis, automation, and isotopomer profiling, and contributed to the production of figures. CS and NB produced illustrations. NB, KN, and CS contributed to writing the manuscript. All authors contributed to review of the manuscript. NB, KN, and JDP acquired funding. All authors read and approved the final manuscript.

## Conflict of interests

The authors declare that they have no conflict of interest.

## Notes

### Competing Interest Statement

The authors have declared no competing interest.

