## Supplemental Figures & Tables for "Isotopically Nonstationary ^13^C Metabolic Flux Analysis in Resting and Activated Human Platelets"

Supplemental Table 1 – Calculated net fluxes
Supplemental Table 2 – Calculated exchange fluxes
Supplemental Table 3 – Comprehensive list of abbreviations
Supplemental Table 4 – MRM ion transitions and MS parameters
Supplemental Table 5 – List of platelet central metabolism reactions and their atom transitions

**FIGURES:**

Supplemental Figure 1 – Simulated labeling distributions for glucose and acetate tracers
Supplemental Figure 2 – Metabolite pool size assessment
Supplemental Figure 3 – Glutamine enrichment of TCA metabolites
Supplemental Figure 4 – Mass isotope abundances and model fits no thrombin case
Supplemental Figure 5 – Mass isotope abundances and model fits plus thrombin case
Supplemental Figure 6 – Aspartate labeling profiles

**TABLES**

**Supplemental Table 1. Calculated net fluxes.**

| Reaction | -thrombin | | | +thrombin | | |
|---|---|---|---|---|---|---|
| | Value | LB95 | UP95 | Value | LB95 | UB95 |
| glucose → g6p | 69 | 49 | 90 | 212 | 186 | 237 |
| g1p ↔ g6p | 23 | 3 | 49 | 178 | 100 | 208 |
| g6p → 6pg | 34 | 22 | 50 | 50 | 24 | 78 |
| 6pg → p5p + co2 | 34 | 22 | 50 | 50 | 24 | 78 |
| p5p + p5p ↔ s7p + gap | 11 | 7 | 17 | 17 | 8 | 26 |
| s7p + gap ↔ f6p + e4p | 11 | 7 | 17 | 17 | 8 | 26 |
| p5p + e4p ↔ f6p + gap | 11 | 7 | 17 | 17 | 8 | 26 |
| g6p ↔ f6p | 58 | 42 | 109 | 340 | 262 | 405 |
| f6p ↔ fbp | 81 | 63 | 132 | 374 | 298 | 426 |
| fbp ↔ dhap + gap | 81 | 63 | 132 | 374 | 298 | 426 |
| dhap ↔ gap | 81 | 63 | 132 | 374 | 298 | 426 |
| gap ↔ 3pg | 173 | 136 | 283 | 764 | 611 | 872 |
| 3pg ↔ pep | 173 | 136 | 283 | 764 | 611 | 872 |
| pep → pyr | 173 | 136 | 283 | 764 | 611 | 872 |
| pyr ↔ lac | 173 | 137 | 208 | 556 | 513 | 599 |
| pyr → aca + co2 | 0 | 0 | 96 | 208 | 59 | 317 |
| acetate → aca | 157 | 26 | 290 | 153 | 48 | 247 |
| aca + oaa → cit | 157 | 26 | 303 | 361 | 104 | 531 |
| cit ↔ akg + co2 | 157 | 26 | 303 | 361 | 104 | 531 |
| akg → succ + co2 | 157 | 26 | 303 | 361 | 104 | 531 |
| succ ↔ fum | 157 | 26 | 303 | 361 | 104 | 531 |
| fum ↔ mal | 157 | 26 | 303 | 361 | 104 | 531 |
| mal ↔ oaa | 157 | 26 | 303 | 361 | 104 | 531 |
| co2 ↔ co2.e | 349 | 71 | 816 | 979 | 326 | 1408 |

**Supplemental Table 2. Calculated exchange fluxes.**

| Reaction | -thrombin | | | +thrombin | | |
|---|---|---|---|---|---|---|
| | Value | LB95 | UP95 | Value | LB95 | UB95 |
| g6p ↔ g1p | 74.43 | 22.75 | 122.24 | $1.9 \times 10^{-6}$ | 0 | 87.86 |
| p5p + p5p ↔ s7p + gap | 34.61 | 17.79 | 62.88 | 40.07 | 18.43 | 68.87 |
| s7p + gap ↔ f6p + e4p | 15.02 | 7.707 | 28.13 | 26.93 | 10.99 | 46.66 |
| p5p + e4p ↔ f6p + gap | $6.0 \times 10^{-6}$ | 0 | 3.208 | 0.0563 | 0 | 9.100 |
| g6p ↔ f6p | $4.4 \times 10^{5}$ | 270.2 | $4.4 \times 10^{5}$ | 5836 | 1305 | Inf |
| f6p ↔ fbp | $8.9 \times 10^{5}$ | 460.5 | Inf | $2.8 \times 10^{5}$ | 1219 | Inf |
| fbp ↔ dhap + gap | 59.36 | 34.59 | 88.42 | 283.83 | 203.2 | 423.48 |
| gap ↔ dhap | 1152 | 385.6 | Inf | 1689 | 897.1 | 5104 |
| gap ↔ 3pg | $6.0 \times 10^{-6}$ | 0 | Inf | $1.0 \times 10^{-7}$ | 0 | Inf |
| 3pg ↔ pep | 5083 | 0 | 5119 | 5296 | 0 | Inf |
| pyr ↔ lac | $1.0 \times 10^{4}$ | 0 | $1.0 \times 10^{4}$ | $1.0 \times 10^{-7}$ | 0 | Inf |
| cit.m ↔ akg + co2 | $1.0 \times 10^{-7}$ | 0 | Inf | $1.9 \times 10^{-6}$ | 0 | Inf |
| succ ↔ fum | $1.0 \times 10^{4}$ | 0 | $1.0 \times 10^{4}$ | $1.6 \times 10^{5}$ | 0 | Inf |
| fum ↔ mal | 9875 | 0 | $1.0 \times 10^{4}$ | $2.0 \times 10^{5}$ | 0 | Inf |
| mal ↔ oaa | $6.0 \times 10^{-6}$ | 0 | Inf | 1546 | 0 | Inf |
| mal ↔ pyr + co2 | $4.0 \times 10^{-6}$ | 0 | 0.5648 | 4.200 | 0 | 14.90 |
| akg ↔ Glu | $2.2 \times 10^{4}$ | 0 | Inf | 5706 | 1529 | Inf |
| Glu ↔ Gln | $1.0 \times 10^{-7}$ | 0 | Inf | $1.0 \times 10^{-7}$ | 0 | Inf |
| oaa + Glu ↔ Asp + akg | 0.1087 | $6.2 \times 10^{-7}$ | Inf | $3.3 \times 10^{-6}$ | 0 | $5.5 \times 10^{-6}$ |
| pyr + Glu ↔ Ala + akg | $5.2 \times 10^{-4}$ | 0 | 0.8567 | $3.7 \times 10^{-5}$ | 0 | 20.07 |
| co2 ↔ co2.e | 9958 | 0 | Inf | $1.0 \times 10^{-7}$ | 0 | Inf |

**Supplemental Table 3. Comprehensive list of abbreviations.**

| Abbreviation | Full Name |
|---|---|
| $^{13}$C-MFA | isotope-assisted metabolic flux analysis |
| 3PG | 3-phosphoglycerate |
| 6PG | 6-phosphogluconate |
| ACA | acetyl-CoA |
| ACD | anticoagulant citrate dextrose |
| AKG | alpha-ketoglutarate |
| ALA | alanine |
| ASP | aspartate |
| CIT | citrate |
| DHAP | dihydroxyacetone phosphate |
| E4P | erythrose 4-phosphate |
| ECAR | extracellular acidification rate |
| F6P | fructose 6-phosphate |
| FBP | fructose 1,6-bisphosphate |
| FUM | fumarate |
| G1P | glucose 1-phosphate |
| G6P | glucose 6-phosphate |
| GAP | glyceraldehyde 3-phosphate |
| GLN | glutamine |
| GLU | glutamate |
| INCA | Isotopomer Network Compartment Analysis |
| INST-MFA | isotopically nonstationary metabolic flux analysis |
| LAC | lactate |
| LC-MS/MS | liquid chromatography tandem mass spectrometry |
| MAL | malate |
| MID | mass isotope distribution |
| MFA | metabolic flux analysis |
| MRM | multiple reaction monitoring |
| OAA | oxaloacetate |
| OCR | oxygen consumption rate |
| P5P | five-carbon sugar |
| PEP | phosphoenolpyruvate |
| PGI2 | prostacyclin |
| PRP | platelet rich plasma |
| PYR | pyruvate |
| S7P | sedoheptulose 7-phosphate |
| SUC | succinate |
| TCA | tricarboxylic acid |
| XF | extracellular flux |

**Supplemental Table 4. MRM ion transitions and MS parameters.** Labeled metabolites were measured by tracking the product ions (Q3) fragmented from each targeted [M-H] parent ion (Q1). Additional listed parameters are as follows: collision energy (CE); collision cell entrance potential (CEP); collision cell exit potential (CXP), declustering potential (DP); and entrance potential (EP).

| Metabolite | Q1 | Q3 | CE | CEP | DP | EP |
|---|---|---|---|---|---|---|
| FBP | 339-345 | 79 | -65 | -14 | -65 | -12 |
| S7P | 289-296 | 79 | -35 | -8 | -65 | -12 |
| 6PG | 275-281 | 97 | -50 | -3 | -20 | -20 |
| G6P | 259-265 | 97 | -50 | -8 | -20 | -10 |
| F6P | 259-265 | 97 | -5 | -14 | -20 | -30 |
| P5P | 229-234 | 79 | -20 | -4 | -50 | -10 |
| 3PG | 185-188 | 79 | -60 | -4 | -60 | -15 |
| GAP | 169-172 | 97 | -50 | -4 | -10 | -6 |
| DHAP | 169-172 | 97 | -20 | -5 | -15 | -4 |
| PEP | 167-170 | 79 | -10 | -8 | -60 | -15 |
| GLU | 146-150 | 102-105 | -20 | -13 | -15 | -10 |
| AKG | 145-150 | 101-105 | -35 | -14 | -15 | -30 |
| GLN | 145-150 | 127-132 | -20 | -2 | -15 | -10 |
| MAL | 133-137 | 115-119 | -35 | -10 | -16 | -5 |
| ASP | 132-136 | 88-91 | -30 | -12 | -15 | -10 |
| SUC | 117-121 | 73-76 | -20 | -12 | -20 | -30 |
| FUM | 115-119 | 71-74 | -70 | -14 | -10 | -30 |
| LAC | 89-92 | 43-45 | -35 | -10 | -25 | -20 |
| ALA | 88-91 | 42-44 | -5 | -2 | -15 | -5 |

**Supplemental Table 5. List of platelet central metabolism reactions and their atom transitions**

| | | |
|---|---|---|
| **Glycolysis** | | |
| glucose (abcdef) | → | g6p (abcdef) |
| g1p (abcdef) | ↔ | g6p (abcdef) |
| g6p (abcdef) | ↔ | f6p (abcdef) |
| f6p (abcdef) | ↔ | fbp (abcdef) |
| fbp (abcef) | ↔ | dhap (cba) + gap (def) |
| gap (abc) | ↔ | dhap (abc) |
| gap (abc) | ↔ | 3pg (abc) |
| 3pg (abc) | ↔ | pep (abc) |
| pep (abc) | → | pyr (abc) |
| pyr (abc) | ↔ | lac (abc) |
| **Pentose Phosphate Pathway** | | |
| g6p (abcdef) | → | 6pg (abcdef) |
| 6pg (abcdef) | → | p5p (bcdef) + co2 (a) |
| p5p (abcde) + p5p (fghij) | ↔ | s7p (abfghij) + gap (cde) |
| s7p (abcdefg) + gap (hij) | ↔ | f6p (abchij) + e4p (defg) |
| p5p (abcde) + e4p (fghi) | ↔ | f6p (abfghi) + gap (cde) |
| **TCA Cycle** | | |
| pyr (abc) | → | aca (bc) + co2 (a) |
| acetate (ab) | → | aca (ab) |
| aca (ab) + oaa (cdef) | → | cit (fedbac) |
| cit (abcdef) | ↔ | akg (abcde) + co2 (f) |
| akg (abcde) | → | succ (bcde) + co2 (a) |
| succ (abcd) | ↔ | fum (abcd) |
| fum (abcd) | ↔ | mal (abcd) |
| mal (abcd) | ↔ | oaa (abcd) |
| mal (abcd) | ↔ | pyr (abc) + co2 (d) |
| **Amino Acid Metabolism** | | |
| akg (abcde) | ↔ | Glu (abcde) |
| Glu (abcde) | ↔ | Gln (abcde) |
| oaa (abcd) + Glu (efghi) | ↔ | Asp (abcd) + akg (efghi) |
| pyr (abc) + Glu (defgh) | ↔ | Ala (abc) + akg (defgh) |

### FIGURES

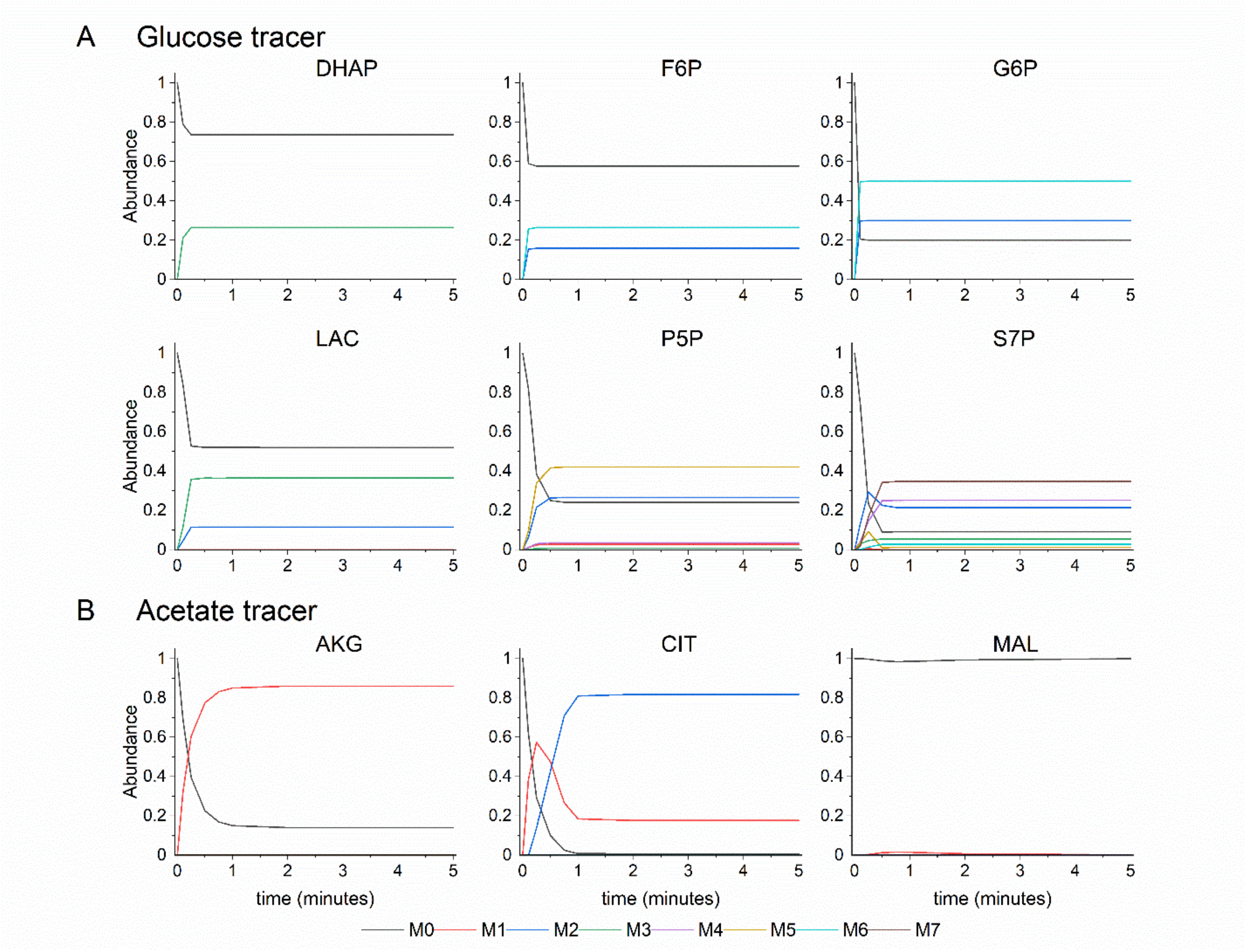


**Supplemental Figure 1. Simulated labeling distributions.** Labeling simulated from (A) 30% [1,2-$^{13}C_2$]glucose, 50% [uniform-$^{13}C_6$]glucose, 20% unlabeled glucose and (B) 25% [1-$^{13}C$]acetate, 25% [2-$^{13}C$]acetate, 50% unlabeled acetate.

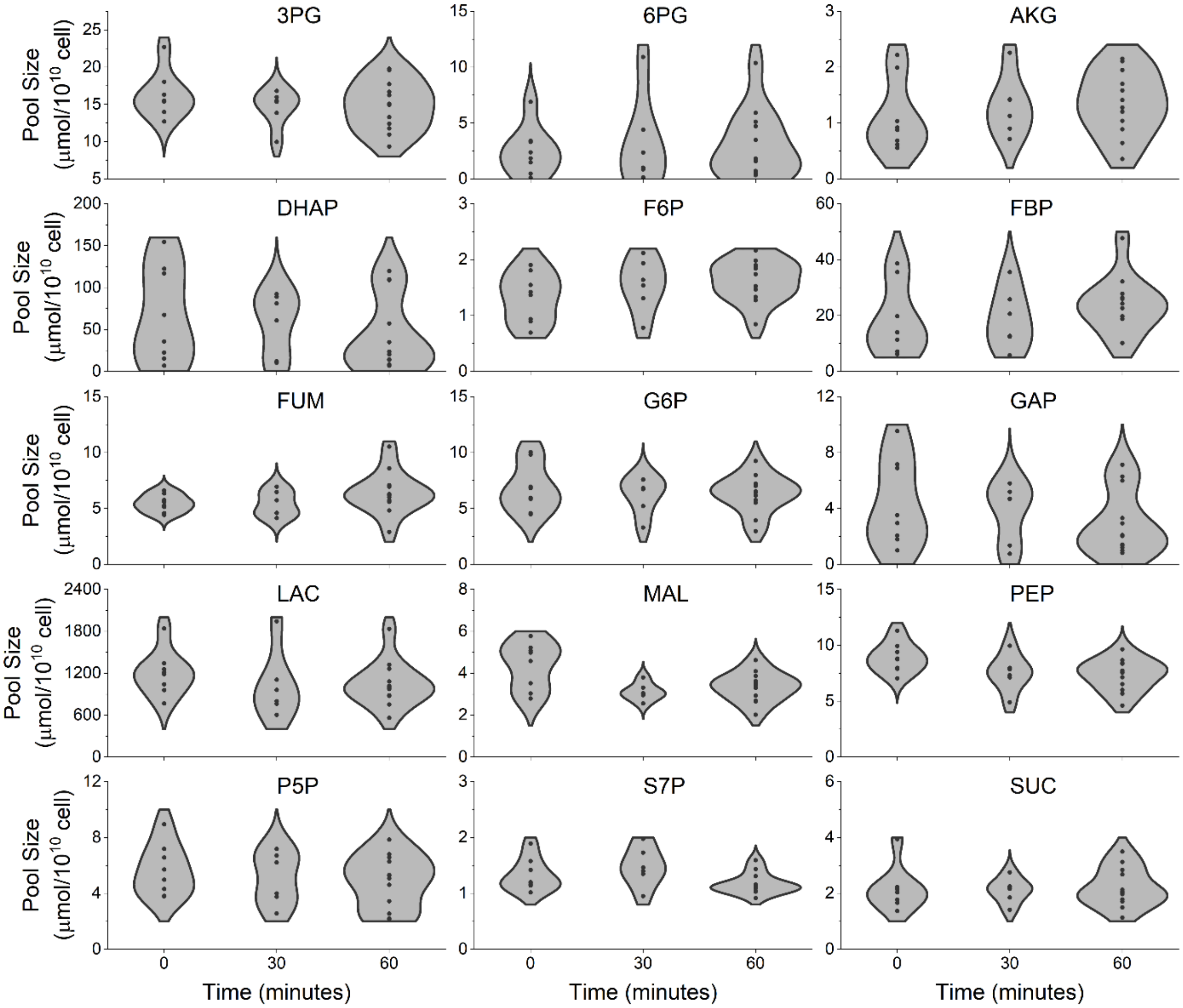


**Supplemental Figure 2. Metabolite pool sizes for platelet central metabolites.** Metabolite pool sizes were measured from washed platelets at 30-minute intervals for one hour. Platelets were allowed to warm to 37°C after washing, prior to the experiment. Measurements were taken with technical replicates (n=3) across adult human female and male donors (n=4). Statistical analysis via ANOVA indicates that metabolite pool size is not determined by time for any metabolite ($p<0.05$).

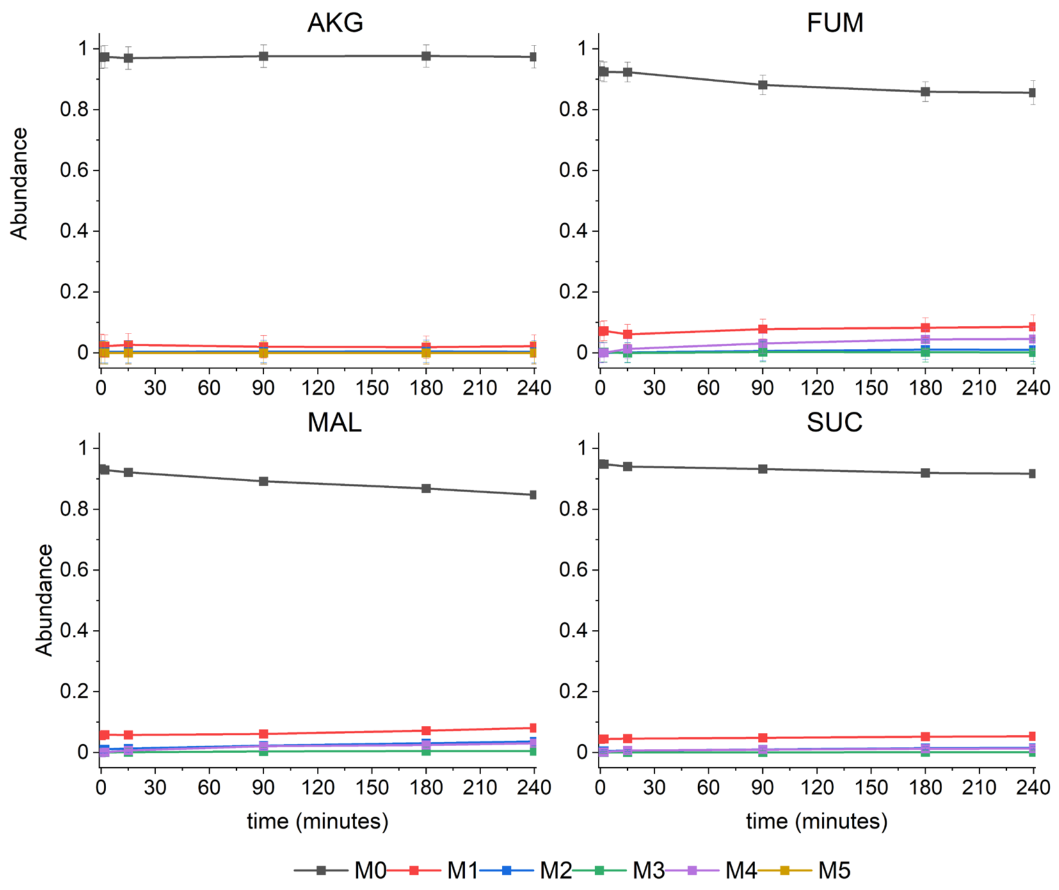


**Supplemental Figure 3. Measured mass isotopomer abundances in TCA metabolites from [U-$^{13}C_5$]glutamine.** Raw mass isotopomer abundances are shown without correction for natural abundance. Error bars represent standard deviation. Lines are not model fits and are included to show data trend.

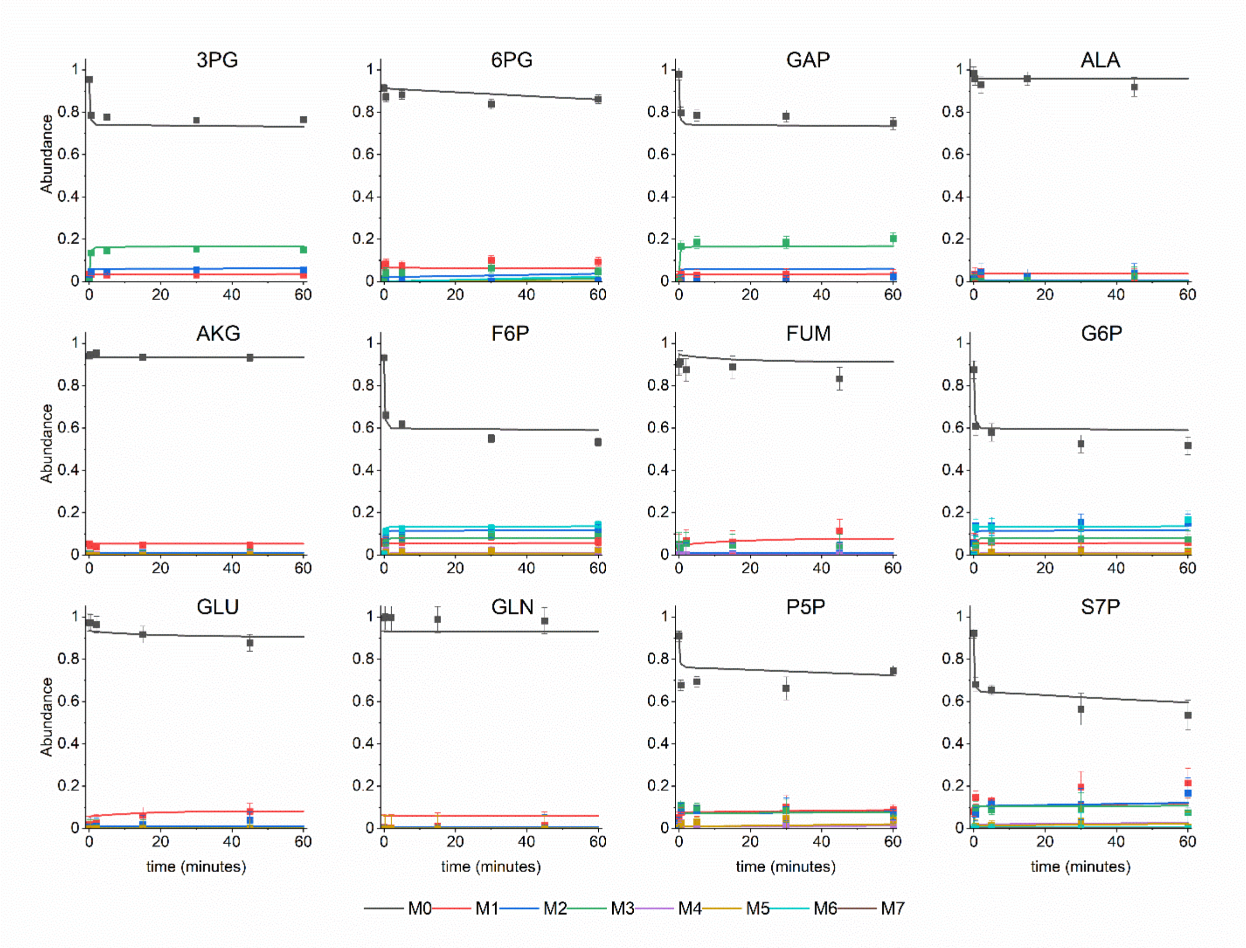


**Supplemental Figure 4. Mass isotopomer abundances (symbols) and model fits (lines) for the no thrombin case.** Labeling for 3PG, 6PG, GAP, F6P, G6P, P5P, and S7P originate from the glucose tracer and labeling for ALA, AKG, FUM, GLU, and GLN come from the acetate tracer. Raw mass isotopomer abundances are shown without correction for natural abundance. Error bars represent standard measurement error.

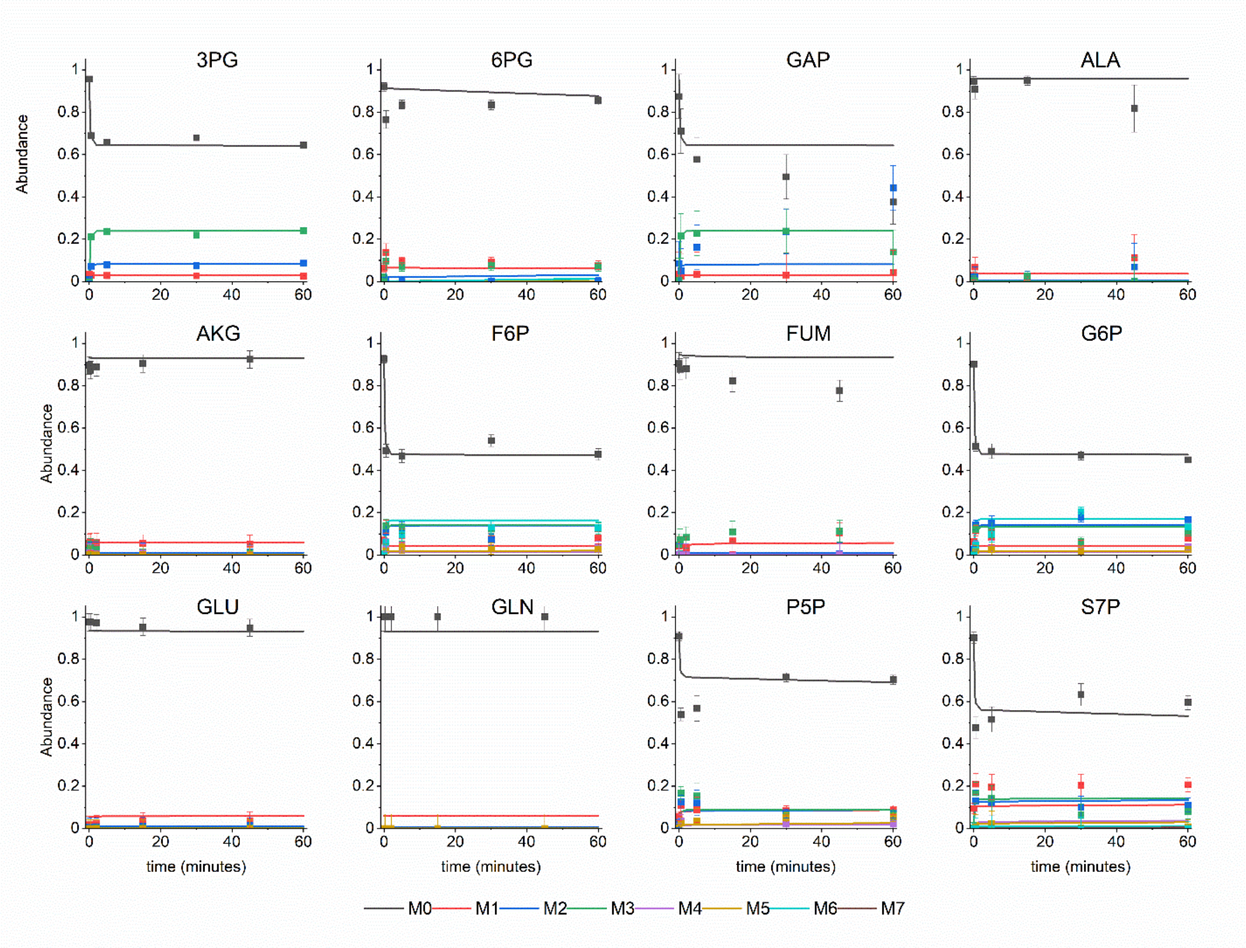


**Supplemental Figure 5. Mass isotopomer abundances (symbols) and model fits (lines) for the 1U/mL thrombin case.** Labeling for 3PG, 6PG, GAP, F6P, G6P, P5P, and S7P originate from the glucose tracer and labeling for ALA, AKG, FUM, GLU, and GLN come from the acetate tracer. Raw mass isotopomer abundances are shown without correction for natural abundance. Error bars represent standard measurement error.

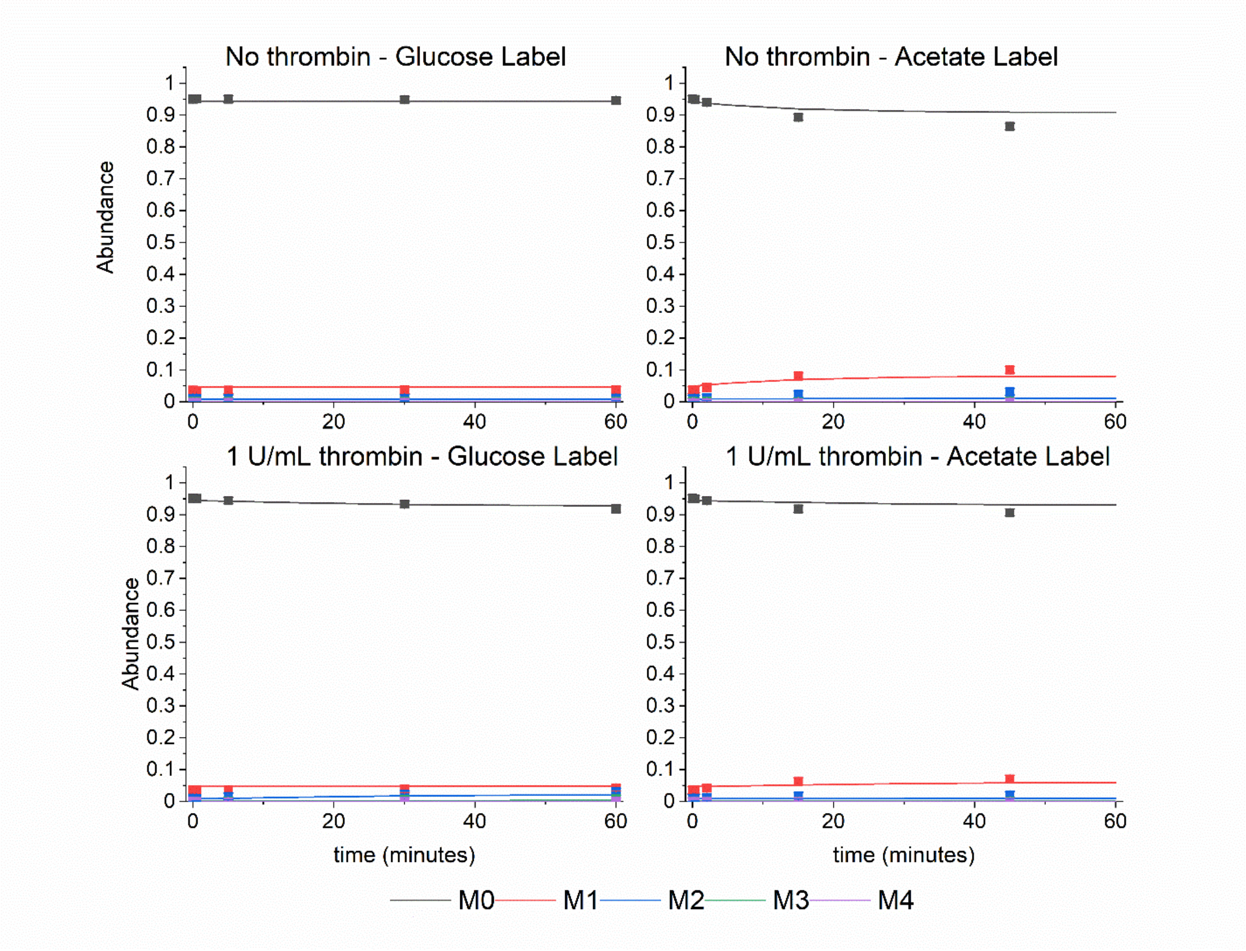


**Supplemental Figure 6. Mass isotopomer abundances (symbols) and model fits (lines) for aspartate.** Raw mass isotopomer abundances are shown without correction for natural abundance. Error bars represent standard measurement error.
